# Reverse Pneumatic Artificial Muscles for Application in Low-Cost Artificial Respirators

**DOI:** 10.1101/2020.05.20.107342

**Authors:** Seyed M Mirvakili, Douglas Sim, Robert Langer

**Author notes:** Dedicated to those who sacrificed their lives in the front line of the combat with COVID-19.

## Abstract

One of the main challenges associated with mechanical ventilators is their limited availability in pandemics and other emergencies. Therefore, there is a great demand for mechanical ventilators to address this issue. In this work, we propose a low-cost, portable, yet high-performance design for a volume-controlled mechanical ventilator. We are employing pneumatic artificial muscles, such as air cylinders, in the reverse mode of operation to achieve mechanical ventilation. The current design of the device can operate in two modes: controlled mode and assisted mode. Unlike most ICU ventilators, our device does not need a high-pressure air pipeline to operate. With the current design, mechanical ventilation for respiration rate ranging from 10 b/min to 30 b/min with a tidal volume range of 150 mL to 1000 mL and I:E ratio of 1:1 to 1:5 can be performed. We achieved a total cost of less $400 USD to make one device. We estimate the device to cost less than $250 USD when produced in larger volumes.

## Introduction

Respiratory dysfunction due to diseases, physical damage to the lungs, or air pollution can be fatal if not treated in time. Artificial ventilation is used to deliver “air” (*e.g.*, oxygen + air/helium/nitric oxide), in a pure form or mixed with drugs, to the lungs, as well as to assist or replace spontaneous breathing. The two major methods of pulmonary ventilation are *manual* insufflation of the lungs (via mouth-to-mouth resuscitation or compressing a bag-valve-mask (BVM)) and *mechanical* ventilation of the lungs with electronically or mechanically controlled equipment. The manual method can be sufficient for temporary cardiopulmonary resuscitation (CPR), while mechanical ventilation is necessary for prolonged pulmonary ventilation. The state-of-the-art mechanical ventilators address the need for all range of respiratory failures; however, their time-to-manufacture, design sophistication, cost, and scalability for deployment in mass emergencies such as pandemics, make them not a viable solution in many situations.

Delivering air with a compliant bladder such as a BVM (*e.g.*, Ambu bag resuscitator) is considered to be among the simplest yet effective method for pulmonary ventilation. While being sufficient for emergency cases, the BVM manual resuscitators cannot be utilized for prolonged ventilation. Moreover, the operation of BVMs requires training and constant attention of the operator. This tedious and repetitive task can be automated to save the time of a well-trained medical staff who can then focus on other aspects of resuscitation. MIT’s E-Vent solution addresses this issue by utilizing an electronically controlled mechanical gripper that periodically presses the compliant bladder of the BVM resuscitator (*1*). Since its introduction, a variety of approaches have been proposed and implemented to automate the operation of BVM resuscitators (*1*). Although automation is achieved, there are several shortcomings associated with interfacing BVM resuscitators with mechanical grippers. Due to the dynamics of deflating the Ambu bag, there can be inconsistencies between breathing cycles. Moreover, it is difficult to monitor the flow rate and volume of air delivered to the lungs due to the compliance of the bag and its geometry. Additionally, excessive and prolong compression/decompression of the bag, if not secured, can lead to material fatigue and leakage which makes it delicate (*2*).

There are two major types of mechanical ventilation: positive pressure ventilation and negative pressure ventilation. In the positive pressure ventilation, a positive pressure of air is applied to the lungs to deliver a controllable volume to the respiratory system. In comparison, in negative pressure ventilation, the chest goes under a negative pressure, which expands the lungs and sucks air into the lungs (*e.g.*, iron lung, chest cuirass). Negative pressure ventilators are considered to be noninvasive.

Positive pressure ventilators were first introduced in the early 1950s to treat polio patients with respiratory paralysis (*3*). They deliver air under a constant volume, pressure, flow rate, or a combination of these parameters. Constant pressure ventilators can be noninvasive or invasive. In noninvasive ventilation, the air is delivered via an interface such as nasal, oronasal, facial masks, mouthpieces, and helmets. Primary modes for noninvasive ventilation are continuous positive airway pressure (CPAP), auto-titrating (adjustable) positive airway pressure (APAP), and bilevel positive airway pressure (BiPAP). Unlike the noninvasive ventilation technique, in the invasive technique, the air is delivered with a tube via endotracheal intubation or tracheostomy. Positive pressure ventilators are the most popular type of respirator. They are used at home (for obstructive sleep apnea), resuscitation in the case of emergency, ICUs, and operation rooms during surgery.

A typical ICU ventilator starts around $2,000 and can be as expensive as $50,000 (*4*). In this work, we are demonstrating a fully functional device from readily available components such pneumatic artificial muscles. Pneumatic artificial muscles are one of the most widely applied actuators in the industry due to their simple design (*5*, *6*). The energy and power density of such artificial muscles do not exceed those of thermal actuators such as shape memory alloy fibers (*7*, *8*) and highly oriented semi-crystalline fibers (*9*, *10*). However, in terms of cycle life and performance stability, PAMs are among the most reliable actuators suitable for precision applications such as in biomedical devices. Air cylinders, one of the sub-categories of artificial muscles, convert pneumatic energy to mechanical energy in a cylinder/piston type structure. We are using this operation mechanism in reverse by applying a linear stroke to the actuator to obtain air pressure/volume to perform the mechanical ventilation.

Thanks to the advances in manufacturing precision glassware, glass syringes of up to 1000 mL capacity are now commercially available. Traditionally, in chemistry labs, glass syringes are used to smoothly deliver an exact volume of a gas to a system or collect a known volume of gas from a reactor. Unlike plastic syringes, the small coefficient of friction between the plunger and the barrel of the glass syringe offers a smooth profile for flow rate and pressure. In our design we are utilizing a 1000 mL glass syringe as the pressure source for mechanical ventilation.

In some anesthesia ventilators, smooth and accurate ventilation is achieved by employing bellows. However, the mechanical work for moving the bellows is provided by the high-pressure (50 psi) central air pipelines of medical centers. Pneumatic-powered mechanical ventilators’ reliance on high-pressure air pipelines makes them less desirable for scenarios where a portable ventilator is needed. Examples of such situations include home care, delivering patients in ambulances, office-based anesthesia practices, and transferring patients from ICU to other units (*e.g.*, imaging, operation room). Moreover, unlike bellows, glass syringes exhibit low compliance. Therefore, accurate tidal volume delivery can be performed.

We believe our proposed solution is reliable for prolonged use and is accurate and low-cost (<$400 USD). Moreover, the device is scalable in emergencies and has a rapid manufacturing time. Our design involves eight major components: 1) stepper motor, 2) motor driver, 3) crank-shaft linkage, 4) 1000 mL glass syringe, 5) power supply, 6) microcontroller, 7) sensor, and 8) interface IO (*i.e.*, pushbuttons, LCD, switches).

The mechanical system is responsible for pushing/pulling the plunger of the glass syringe and is digitally controlled by the microcontroller (Figure 1A). The outlet of the glass syringe is connected to the breathing circuit (Figure 2A). The primary role of the breathing circuit is to enable the syringe to take in air from a reservoir bag or the ambient environment when the glass syringe is pulling. The air is then delivered to the lungs when the glass syringe is pushing without escaping from the intake pathway (Figure 2B). In the current design, we are using a mechanical PEEP valve, as shown in Figure 2B. It is possible to use a digitally controlled PEEP valve. The user interface lets the operator set parameters, stop/start the device, and monitor the pressure and flow rate on the LCD (Figure 1C).

**Figure 1.**
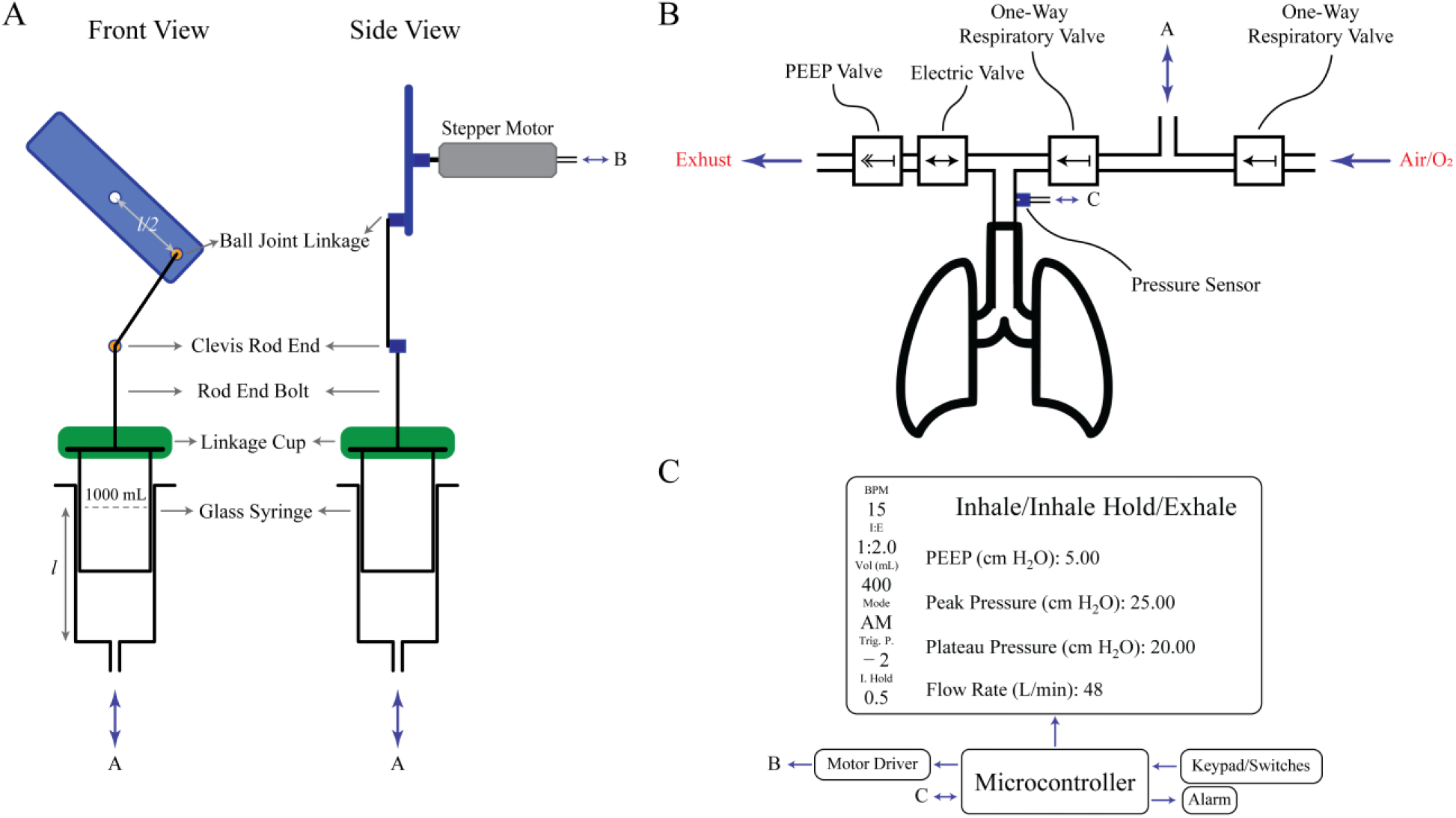
High-level schematic of the ventilator. (A) Illustration of the mechanical system performing the ventilation. (B) Schematic of the breathing circuit interfaced with the ventilator and the lungs. (C) High-level schematic of the electrical components used in the mechanical ventilators and its user interface.

**Figure 2.**
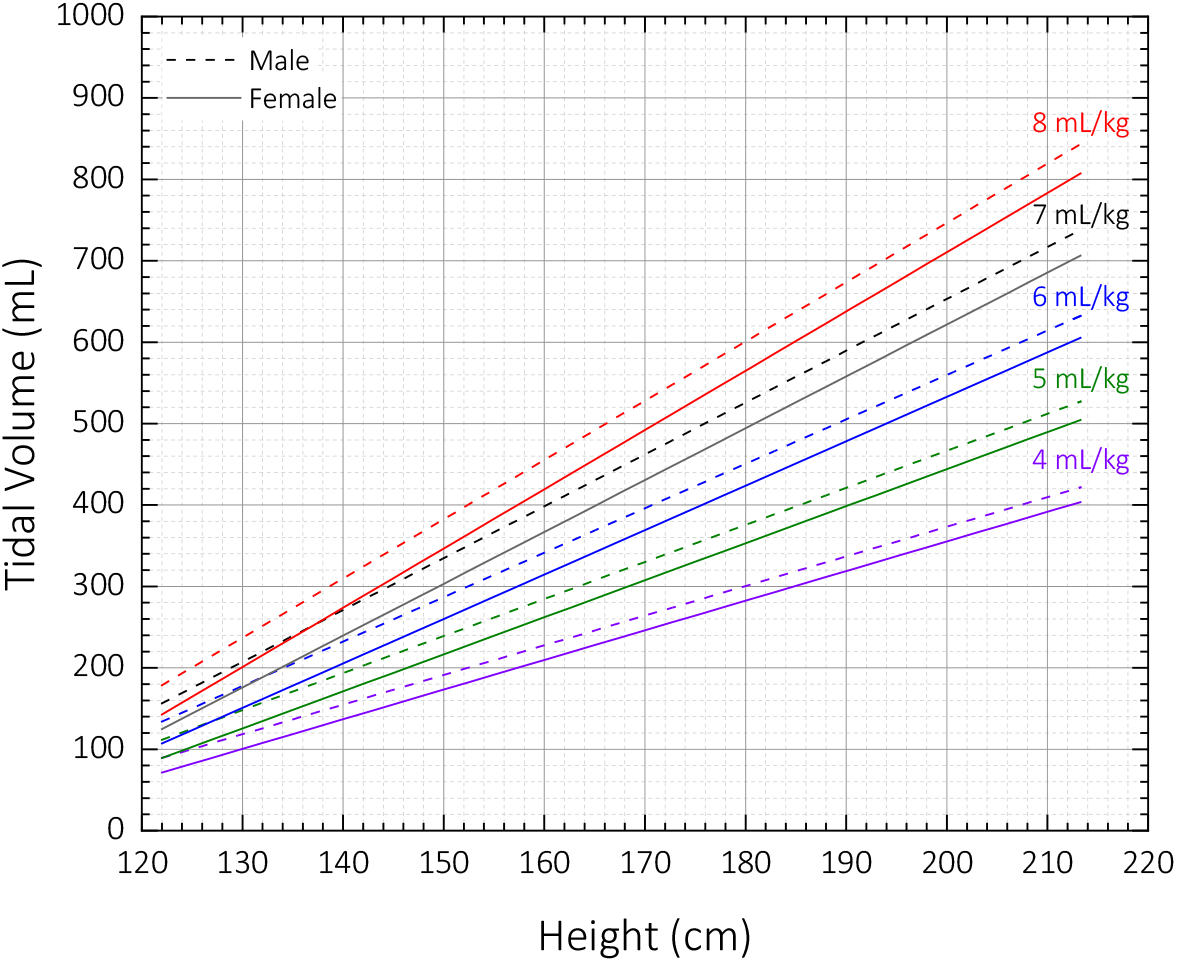
Tidal volume as a function of body height for different gravimetric tidal volumes for male and female patients.

The plunger of the glass syringe is linked to the crank by two joint linkages. The plunger is first attached to a rod end bolt by a resin (materials and methods). The rod end bolt is connected to a clevis rod end, which can freely rotate by 180°. The clevis rod end is linked to the crank via a threaded rod and a ball joint linkage. Finally, the crank is attached to the stepper motor via a flange-mount shaft collar. The crank-shaft linkage converts the rotational motion of the stepper motor into a linear motion. The dynamics of the movement can be controlled by applying different pulse functions to the stepper motor (materials and methods).

The outlet of the syringe is connected to a breathing circuit that connects to the lungs. We are employing the breathing circuit from an Ambu bag for two reasons. First, the one-way respiratory valves, the positive end expiration pressure (PEEP) valve, and the pressure gauge valve are compact and easy to integrate with our design. Second, in the case of failure, the Ambu bag can be operated manually to resuscitate the patient in place of the device.

Our proposed device is categorized as an ACV (assist-control ventilator) device, previously known as the continuous mandatory ventilation (CMV). The device controls the volume and can operate in two modes. The first mode is the *controlled mode* (*e.g.*, CM mode), which is suitable for sedated and paralyzed patients. The second mode is the *assisted mode* (*e.g.*, AM mode) for assisting patients who can breathe spontaneously but need extra volumes of air or higher FiO_2_ (*i.e.*, the fraction of inspired oxygen). Due to the urgent need for ventilators in the current pandemic (COVID-19), we are only demonstrating the mentioned two modes. More modes of operation can be introduced on the same hardware by adding more program logic. The key parameters that control the function of ventilators include:

*Tidal Volume (VT)* is the volume of air displaced between inhalation and exhalation when extra effort is not applied. The gravimetric tidal volume is often reported in the units of cc or mL per predicted body weight (kg). Typical values for patients with acute respiratory distress syndrome (ARDS) are between 4 – 8 mL/kg. The predicted body weight (PBW) for male and female patients can be estimated from the height of the patients according to the ideal body weight equation described in the following equations (*11*):

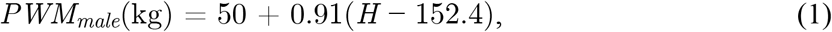

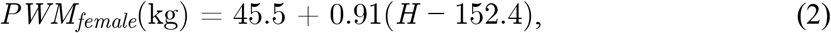

where *H* is the height measured in unit of cm. Figure 2 illustrates the tidal volume as a function of the patient’s height for different gravimetric tidal volumes. As shown in Figure 2, the range for tidal volume is between 70 mL to 840 mL.

*Respiratory Rate (RR)*, also known as breath per minute (BPM), is the number of respiration cycles that occur per minute. The typical range for an adult is between 12 – 20 b/min (*12*). Respiratory rate of up to 30 b/min is also reported for COVID-19 patients (*1*, *13*).

*Inhalation/Exhalation ratio (I*:*E)* defines the time ratio between the inhale and the exhale cycle. The typical value for a healthy human is 1:2 and is reduced to 1:4 or 1:5 in the presence of obstructive airway disease (*14*).

*End-inspiratory hold* is defined as the hold time occurring at the end of the inspiration cycle. The end-inspiratory hold maneuver is performed to eliminate the pressure contribution from the airway resistance and reveal the pressure in the alveoli (*15*). In the current design, the value is set as the percentage of the inhale time.

*Inhale trigger pressure* is only applied in the assisted mode. This represents the slight negative pressure that is developed in the breathing circuit when the body attempts to inhale. By adjusting this parameter, we can ensure the device is synchronized with the spontaneous respiration rate of the patient.

*Positive end-expiratory pressure (PEEP)* is the gauge pressure in the lungs (alveolar pressure) developed at the end of expiration. The PEEP is typically applied by the ventilator at the end of each breath to reduce the likelihood of the alveoli to collapse. This positive pressure ‘recruits’ the closed alveoli in the sick lung and improves oxygenation. In this we work, we employed a mechanical PEEP valve.

Depending on the condition of the patient (*e.g.*, anesthesia, oxygen therapy, obstructive sleep apnea, SARS), different combinations of parameters are used. For example, for acute respiratory distress syndrome (ARDS), which also occurs with COVID-19, a tidal volume of 400 mL to 600 mL with an *I*:*E* ratio of 1:2 to 1:4 with respiration rate of 12 b/min to 16 b/min is used (*13*).

We made a test setup to evaluate the performance of the design. As depicted in Figure 3, the setup is made of readily available components. To accelerate the prototyping of the setup, we built the breathing circuit from plastic tubes and tubing adaptors from fast shipping vendors. It is easy to replace the current breathing circuit with low-cost FDA approved tubing with close no modification of the device.

**Figure 3.**
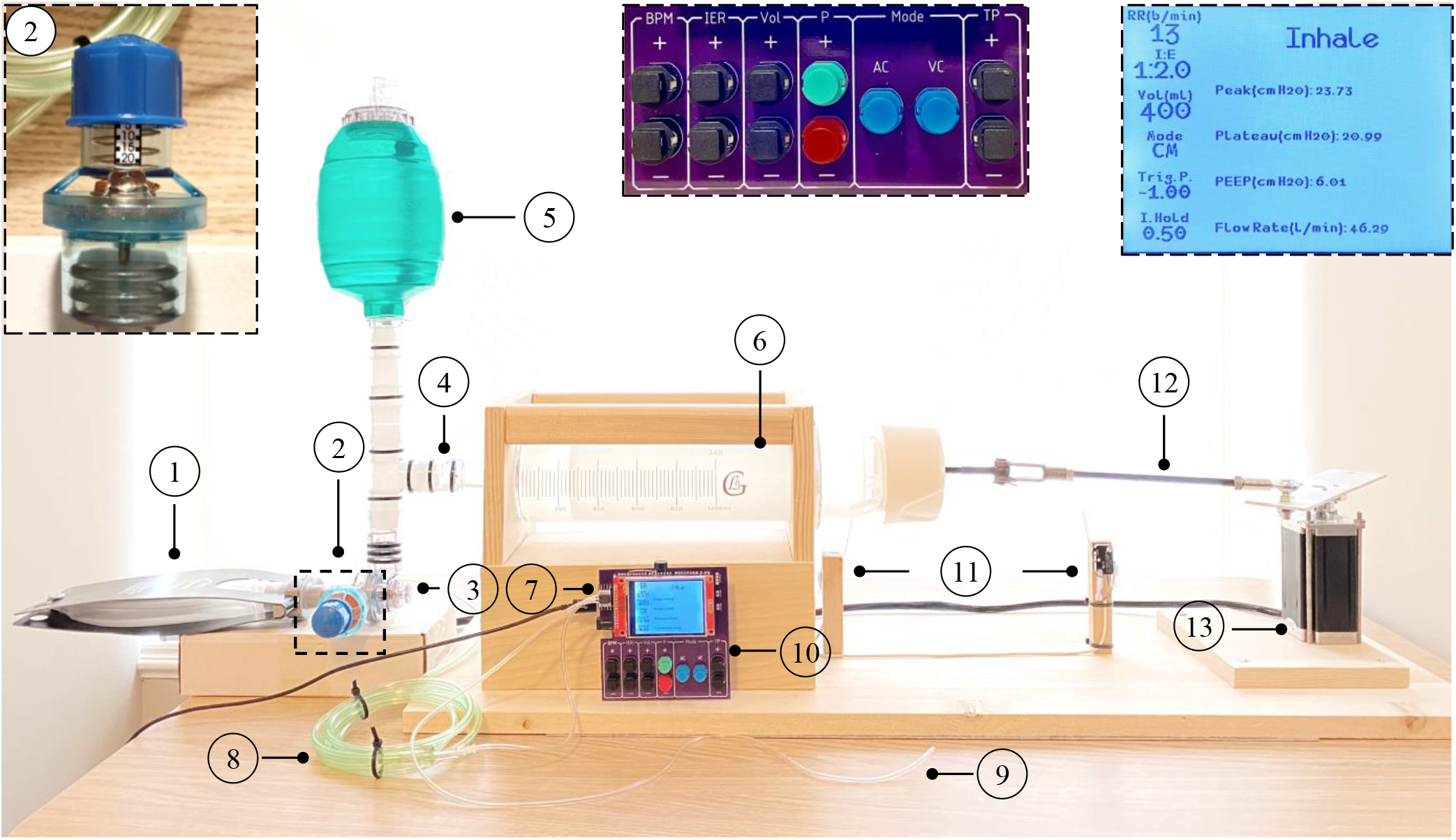
Image of the setup. The components used in this design are: (1) a 1000 mL test lung (compliance: 25 mL/mbar at a tidal volume of 500 mL and PEEP of 0 mbar. Resistance: 20 mbar/L/s), (2) PEEP valve – adjustable from 0 to 20 cmH_2_O, (3) mask adaptor of the BVM resuscitator, (4) tubing and tubing adaptors, (5) resuscitation bag of the BVM with the one-way intake valve. The reservoir bag for oxygen connects to the top of the BVM bag and is not shown here, (6) 1000 mL BLG glass syringe, (7) differential pressure sensor, (8) tubing connecting one port of the pressure sensor to the mask adaptor, (9) tubing connected to the free port of the pressure sensor. This tube was only used in the assist mode to simulate inspiration effort to trigger the device. (10) controller with the keypad and display, (11) two limit switches, (12) crank-shaft linkage (materials and methods), and (13) the stepper motor. Insets: top-right is a zoomed-in snapshot of the display during ventilation and top middle is a zoom-in image of the keypad for the current design. The green button starts the ventilation process and the stop button stops. BPM, IER, Vol, Mode, and TP are the buttons to set the respiration rate, *I*:*E* ratio, tidal volume, mode of operation, and the trigger pressure.

For logging data, we used a sampling rate of 50 ms. For low respiration rates, 50 ms sampling is sufficient. However, for higher respiration rates (> 25 b/min), to log the data accurately faster sampling rate is required, which comes at the expense of stressing the microcontroller.

In volume-controlled mode, the ventilator does not sense any efforts from the patient for breathing. It provides the tidal volume at a specific respiration rate and the *I*:*E* ratio that is configured by the operator. Our device can operate within the range of the parameters listed in Table 1.

**Table 1.**
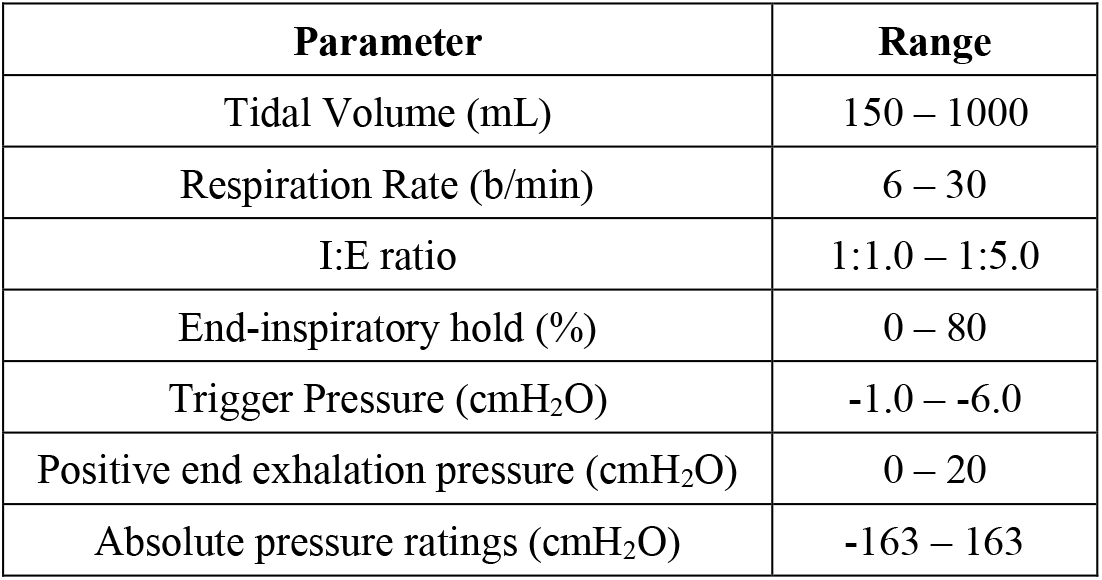
The ventilator device ratings.

The pressure is monitored through the whole respiration cycle; however, the tidal volume and flow rate are calculated from the position of the stepper motor. Therefore, the volume and the flow rate numbers are representative of the inspiration cycle only and data for the expiration phase on the plot represent those of the glass syringe. During the expiration cycle, the patient respiratory system is decoupled from the glass syringe and it is performed independently by the patient via the mechanical recoil of the lungs. To be able to record the patient’s volume and flow rate during the exhale cycle, extra peripherals are needed.

To evaluate the performance of the device, we swept the parameters, tested edge cases, and characterized boundary conditions that are used in real life scenarios as well. Figure 4 illustrates the volume, pressure, and flow rate for nine test cases (tabulated data in supporting information). We performed the end-inspiratory hold maneuver with a hold time of 50% to obtain the plateau pressure for each case. For experiments Figure 4A, we kept the *I*:*E* ratio constant at 1:2 but increased the *RR* from 12 b/min to 30 b/min while decreasing the *VT* from 750 mL to 300 mL. As we decreased the tidal volume, both the peak pressure and plateau pressure decreased as well. This correlation can be explained by the fact that less air is moved to the test lung; therefore, a smaller pressure is developed inside it. For experiments in Figure 4B, we kept the *VT* at 500 mL and *RR* at 13 b/min but increased the *I*:*E* ratio from 1:1 to 1:5. In this case, since the volume is kept constant. Therefore, the elastic component of the peak pressure (*P_plateau_* – *PEEP*) is also constant. However, the resistive component of the peak pressure increases (*P_peak_* – *P_plateau_*). This increase can explain by the fact that the resistive pressure is a function flow rate (supporting information). As illustrated in Figure 4, the cycles are very consistent and repeatable. Any slight discrepancies in the data are most likely due to the under-sampling of the parameters.

**Figure 4.**
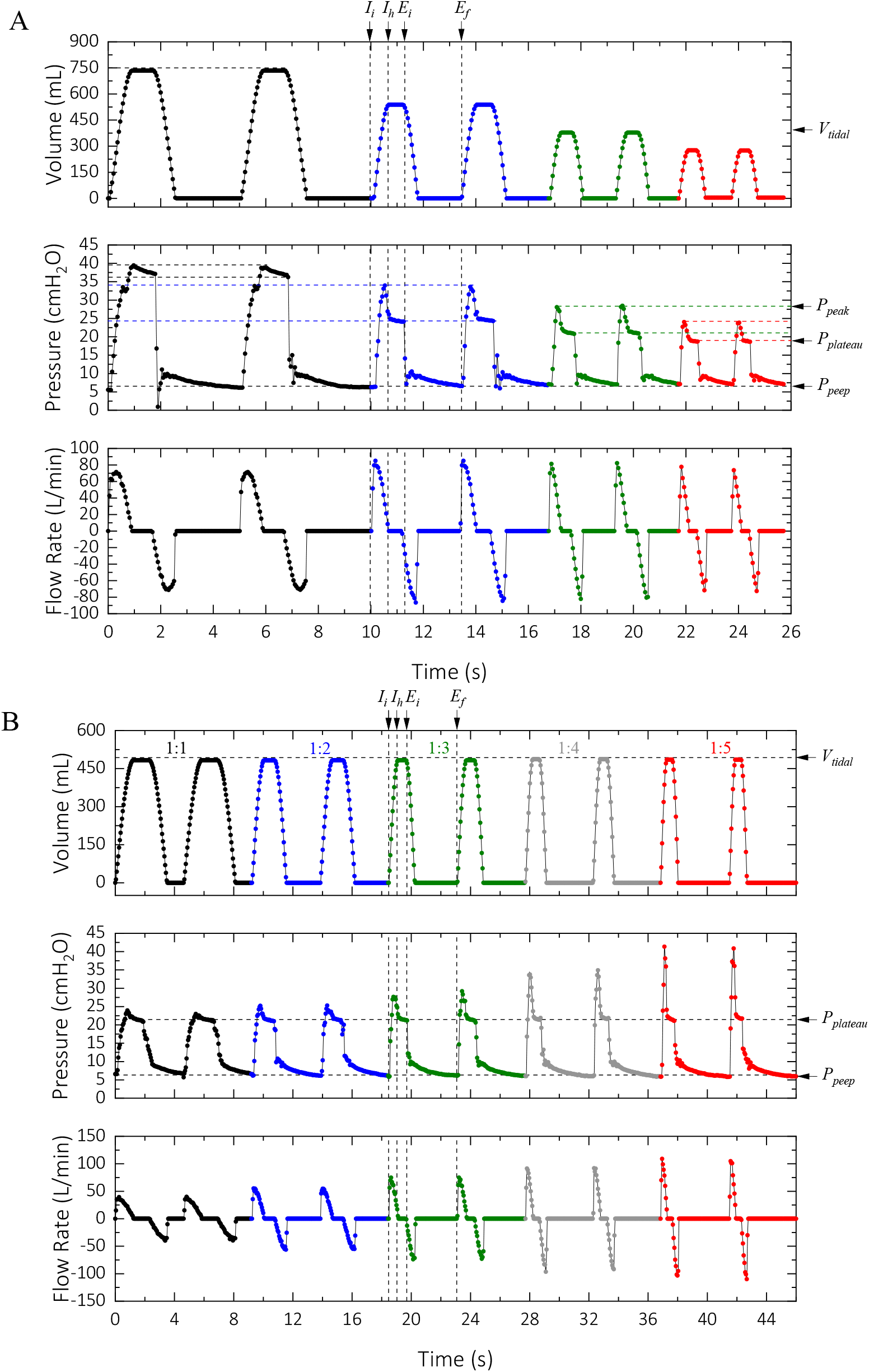
Performance of the ventilator in the controlled mode with a different set of operating parameters. Setting parameters: (A) Black: *VT* = 750 mL, *RR* = 12 b/min; blue: *VT* = 550 mL, *RR* = 18 b/min; green: *VT* = 400 mL, *RR* = 24 b/min; red: *VT* = 200 mL, RR = 30 b/min. The *I*:*E* ratio and the PEEP were set to 1:2 and 6 cmH_2_O for all the cases. (B) The device setting for this case was *VT* = 500 mL, *RR* = 13 b/min with *I*:*E* ratio of 1:1 (black), 1:2 (blue), 1:3 (green), 1:4 (gray), and 1:5 (red). The PEEP was set at 6 cmH_2_O for all cases.

The breathing cycle for the patient consists of three major cycles. The inhale cycle which starts at *Ii*, the inhale hold cycle which starts at *I_h_*, and the exhale cycle which starts at the end of inhale cycle (*E_i_*) and ends at *E_f_*. The exhale cycle for the glass syringe/motor includes one more cycle which is the exhale hold – the bottom flat region on the volume plot in Figure 4. In the exhale hold cycle, the plunger stays in that position (corresponding to the set tidal volume) until the next inhale cycle is triggered internally or by the patient’s attempt to inhale.

To better evaluate the performance of our device, we performed ventilation of a test lung under identical device configurations used for a COVID-19 patient with a Maquet Servo-I ventilator (Figure 5). Figure 5 shows the results for a configuration of 400 mL delivery at *I*:*E* ratio of 1:1.7, respiration rate of 16 b/min, and PEEP of 5 cmH_2_O. This configuration resulted in a peak pressure of 22 cmH_2_O with a plateau pressure of 20 cmH_2_O with the ventilator. The slight discrepancies are due to the small differences in the mechanical compliance of the test lung/the breathing circuit that we used and the patient’s lung/the breathing circuit that was used with the Maquet Servo-I ventilator. To test reproducibility of our device, we logged the performance parameters for 100 consecutive cycles for this configuration with inspiration hold time of 50% (0.7 s). As Figure 6A illustrates, the performance parameters are very consistent over the entire duration. We measured mean peak pressure of 22.20 ± 0.22 cmH_2_O, mean plateau pressure of 20.50 ± 0.10 cmH_2_O, and mean PEEP of 4.98 ± 0.11 cmH_2_O. We performed a long cycle life test under VT = 500 mL, I:E = 1:2, RR = 12 b/min, and inspiration hold = 50%. We measured mean peak pressure of 23.98 ± 0.37 cmH_2_O, mean plateau pressure of 21.22 ± 0.36 cmH_2_O, and mean PEEP of 5.41 ± 0.21 cmH_2_O (Figure 6B). The slight drift in the PEEP value is due the mechanical nature of the PEEP valve that we used. This drift increased the peak and plateau pressure as well.

**Figure 5.**
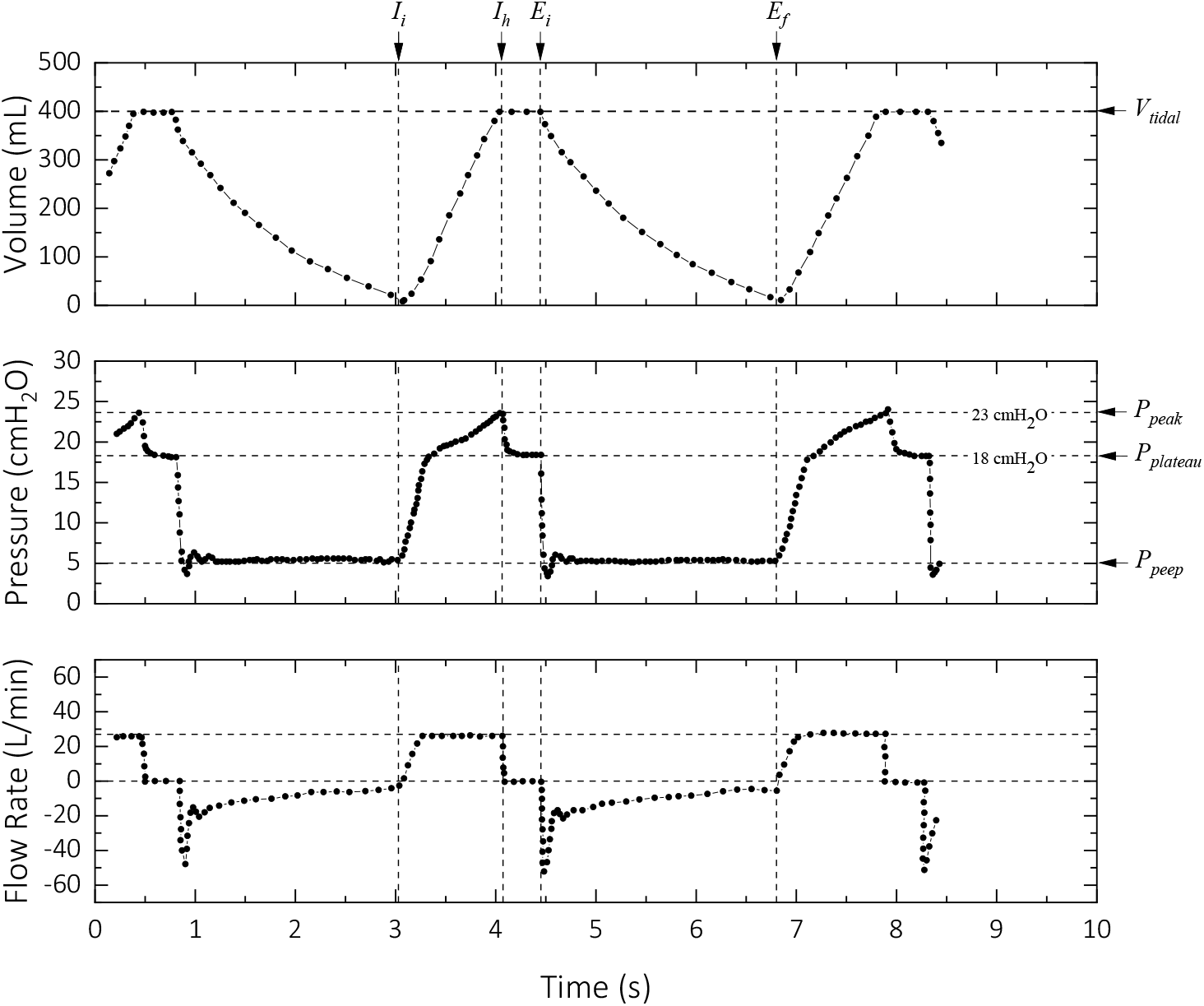
Performance of a Maquet Servo-I Ventilator configured to deliver a tidal volume of 400 mL with PEEP of 5 cmH_2_O and *I*:*E* ratio of 1:1.7. The plot is regenerated by digitizing an image from the screen of the device under operation for a COVID-19 patient. Due to the smoothness of the volume and flow rate profiles, a lower spatial sampling rate was used for the digitization.

**Figure 6.**
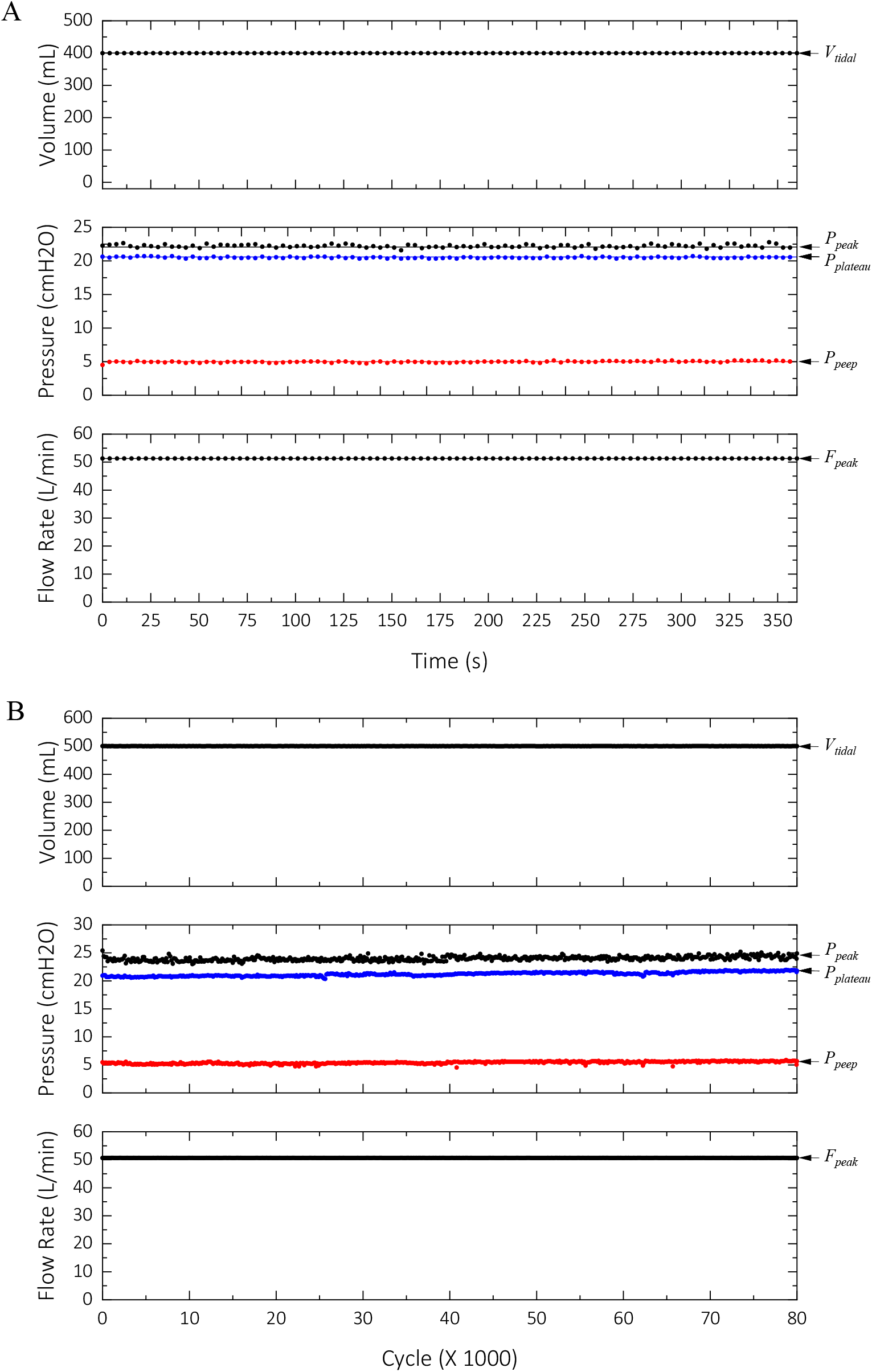
(A) Performance of the device for 100 cycles. The test configuration was *VT* = 400 mL, *I*:*E* = 1:1.7, *RR* = 16 b/min, with inspiration hold time of 50%. As shown, the timing of the cycles is precise. (B) Performance of the device over 80,000 cycles. The test configuration was *VT* = 500 mL, *I*:*E* = 1:2, *RR* = 12 b/min, with inspiration hold time of 50%. The slight drift in the PEEP value increased the peak and plateau pressure values as well.

It is important to note that due to the kinematics of the actuator (supporting information), the plunger of the glass syringe does not travel linearly but rather exhibits a sinusoidal profile (supporting information). Therefore, the flow rate has a ‘sinusoidal’ characteristic. However, it is possible to digitally control the stepper motor to obtain a linear profile for the tidal volume, similar to that of Figure 5.

The tidal volume and flow rate were almost constant, which is due to the fact that in every cycle, the stepper motor’s position was recalibrated with the limit switch. Therefore, there was no error, drift, or inconsistencies between successive cycles.

Modern ventilators deliver oxygen (mixed with air) to the lungs and monitor the CO_2_ during expiration. Our device does not have a capnograph internally; however, an external capnograph can be used to monitor the CO_2_. The reservoir bag of the BVM that we employed in our design can be filled with oxygen to increase the FiO_2_. Moreover, anesthetic gas agents (*e.g.*, sevoflurane, isoflurane) can be delivered via the medication port of the BVM mask adaptor for patients under anesthesia.

Similar to the controlled mode, in the assisted mode a constant volume of air is delivered to the lungs. However, in this mode, the inspiratory cycle is triggered by the patient’s attempt to inhale. The effort to inhale generates a slight negative pressure in the respiratory system which is used to trigger the device. Depending on the condition of the patient, a trigger pressure between −1 to −5 cmH_2_O is configured for this mode of operation. As a safety feature, if the inhale attempt is not detected within a fixed period (set by the respiration rate and the *I*:*E* ratio), the device automatically defaults to the CM mode and notifies the operator via an alarm. To test this mode, we used 500 mL for the tidal volume, a respiration rate of 16 b/min with an *I*:*E* ratio of 1:2, and trigger pressure of −1 cmH_2_O. To simulate the negative pressure generated in the attempt for inspiration by the lungs, we applied slight positive pressure to the open end of the differential pressure sensor (Figure 3). As depicted in Figure 7, when pressure drops below −2 cmH_2_O, the device starts the ventilation and delivers the set tidal volume. In the third cycle of Figure 7, we made no ‘attempt’ for inhale and the device defaulted to the controlled mode and operated continuously as expected (Figure 7).

**Figure 7.**
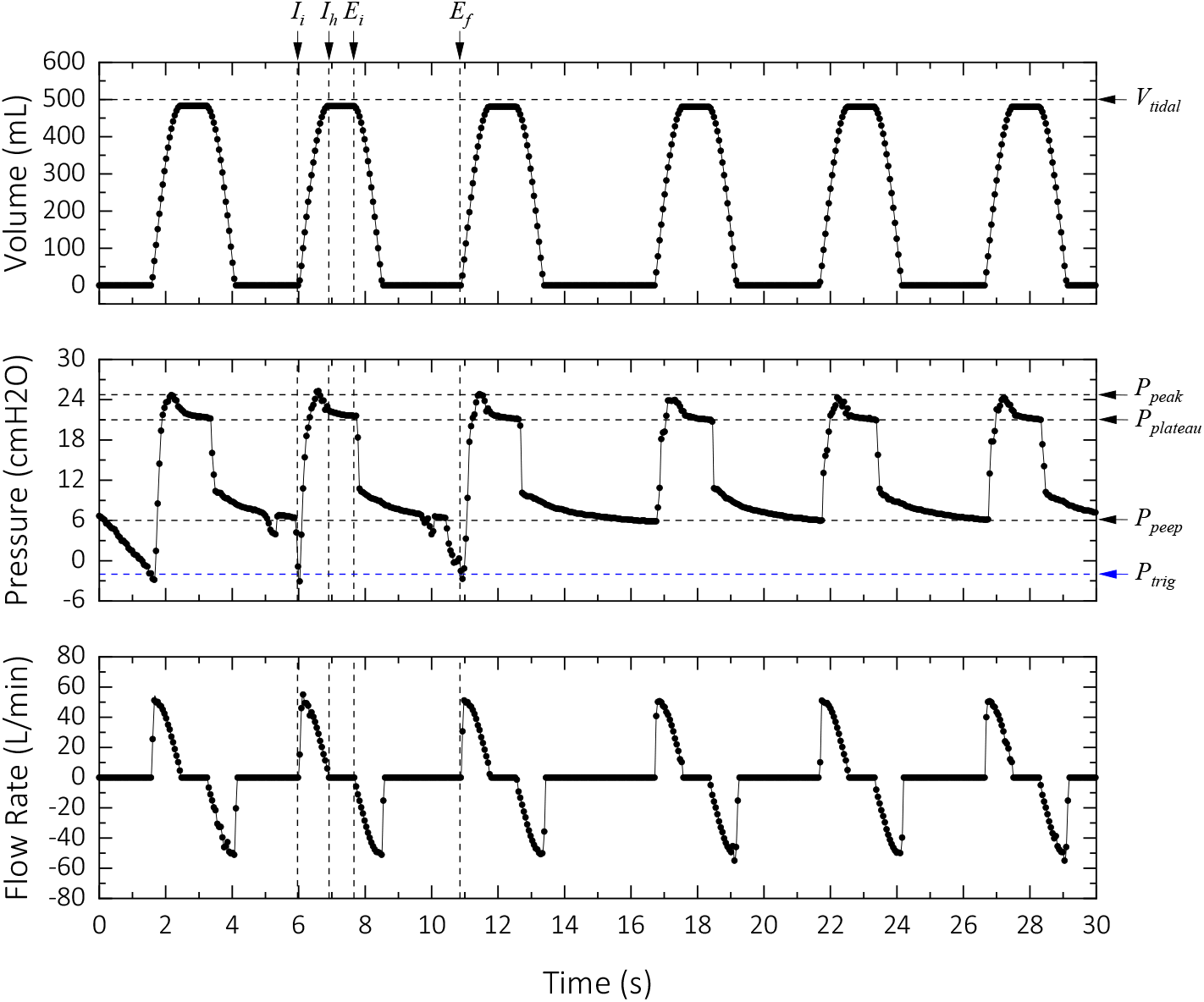
Assisted mode test results. The device was configured to deliver 500 mL at a respiration rate of 12 b/min with an *I*:*E* ratio of 1:2.

Our design rationale is based on optimizing the performance (*e.g.*, accuracy, portability, cycle life), simplicity, time to manufacture, scalability, and cost while utilizing generic and readily available components. Unlike pneumatically powered ventilators (*e.g.*, Bird Mark 7, Bio-Med MVP-10, Servo-I), the driving force in our proposed device is provided by a robust high precision stepper motor, which makes it portable and reliable for prolonged use. To simplify the manufacturing procedure, we avoided 3D any of the components and used readily available materials (Figure 2). We achieved a total bill of materials cost of less than $400 USD for making the prototype (materials and methods).

Our proposed device does not introduce any new modes of operation for ventilators. In other words, standard parameters and modes that are available on any ventilator are used in our solution. Therefore, the adoption of this device could be almost instantaneous. We anticipate most medically trained operators should be able to operate this without any further training or instructions. In the current case of public health emergency, governmental health organizations have relaxed the approval process for mechanical ventilators. For example, Health Canada stated “a summary of clinical evidence, including clinical study results and/or literature review, in accordance with section 4(1)(g) (*i.e.*, the known information in relation to the quality, safety and effectiveness of the device) should be provided if and only if the ventilator includes novel features and concepts. Otherwise, the ventilator is an established technology and clinical study data is not required” (*16*, *17*).

In designing this device, we took into consideration the safety guidelines for designing a ventilator and incorporated multiple safety features in our design. These measures include:

- The pressure is constantly monitored if it passes certain level, that are set in the software, the alarm is triggered to notifies the operator. For the first level (currently set at 20 cmH_2_O), the alarm sounds periodically, and the device continues to function. For the second level (currently set at 40 cmH_2_O), the alarm sounds continuously and halts the machine mid-operation. A mechanical safety valve is used to release the pressure should this situation occur.
- In the assisted mode, if the device does not detect an attempt for inspiration from the patient, it automatically switches to the controlled mode and operates according to the set values.
- In the event of an electronic failure, by default the stepper motor turns off. This feature enables the operator to manually actuate the glass syringe easily from the crank joint. In this case, the tidal volume can be read from the measurement markings on the glass syringe.
- In the event to of a mechanical failure, the operator can immediately disconnect the device from the breathing circuit and use the already attached BVM to perform the ventilation manually.
- The tidal volume is limited by two limit switches. The limit switches change the motor’s direction of rotation when toggled. One of them is used to recalibrate the position of the motor in every breathing cycle. Moreover, the limit switches prevent the motor from going past a certain angle, which could potentially damage the device.
- The disassembly process of the device for maintenance and cleaning is very straightforward. Moreover, glass is relatively chemically non-reactive and resistant to disinfectants.

Historically, to make a ventilator there is a remarkably high barrier of entry due to its targeted field being very niche and specific (*i.e.*, medical field). The high level of regulations and requirements to make and sell such medical devices adds further to this entry barrier. We are achieving a potential advantage by substantially lowering the barrier of entry to manufacturing, deploying, and owning a mechanical ventilator. The COVID-19 pandemic has brought to light the need for simpler and readily available devices that can address the urgency and impact of the situation rather than only relying on high-end all-rounded “perfect” devices.

We focused our fundamental design on addressing the demand for ventilators in life-death situations. Therefore, only the core and essential features, such as the safety features mentioned, as well as the two modes of operation are implemented. Contributing to the cost of high-end ventilators have are the additional quality of life improvements and features which come with the device. Common examples include capnograph, blood oximeter, additional storage for data logging/analysis, larger and more sophisticated LCD interface, IP rating, and IK impact resistance rating. We avoided using custom made parts (*e.g.*, 3D printed components, machined gears, injection-molded pieces) and utilized off-the-shelf components to cut down the cost further. For example, the precision machining of rigid materials such as Teflon or metals for making a 1000 mL cylinder/piston is a subtractive manufacturing method that is time-consuming, labor-intensive, and expensive.

The demand for this device in underdeveloped countries will be much higher due to the sheer cost of high-end first-class ventilators. Moreover, underdeveloped countries typically have a higher population density, which, in turn, creates a higher demand for the number of medical devices. In emergencies such as pandemics where the need for ventilators overwhelms the number of ventilators available, our device can act as a temporary or substitute to high-end medical grade ventilators. Moreover, these ventilators could be part of the first aid or medical kit in large companies or institutions such as universities and airports. Another possible application would be in education and training institutions where hands-on experience and time with such devices not only required but beneficial. Considering precision for the tidal volume delivery that we achieved, we anticipate that our proposed ventilator can be used as an anesthesia ventilator as well.

Future work on this device can be divided into two major areas: improving the device hardware and expanding functionality. Currently, a generic Arduino microcontroller is being used due to its availability and community support, making it ideal for prototyping. For optimization purposes, we could move away from Arduino to a more specialized microcontroller or an SoC (System on Chip). A more feature specific microcontroller unit would lay the foundation for more functionality, such as real-time data analysis, remote monitoring with WiFi, and more compact circuitry. Another significant component that could be improved on is the user interface (UI) and user experience (UX) of the device as well as more sensors (*e.g.*, flow sensor, temperature sensor, EKG, blood oximeter, capnograph). An example of improving the UI/UX of the device would be to add a real-time plot and display more parameters on a larger display similar to high-end ventilators.

The other major area for improvement is the functionality of the device. Currently, a contact volume ventilation is possible via only CM and AM modes with our device. Constant pressure and constant flow rate ventilation with more modes can be incorporated with a minimum change to the hardware architecture. Moreover, a servo-controlled heater can be attached to the body of the syringe to control the temperature of the gas before delivery. A more compact design can be implemented by utilizing a linear voice coil actuator and equip the software with a system identification technique to measure the lung’s compliance and other mechanical properties (*e.g.*, viscoelastic behavior) in situ.

Finally, long-term (weeks) porcine trials should be run to validate the usage of the device for prolonged ventilation. The clinical trials can then be performed to increase the confidence in our device in the medical field.

## Conclusion

In this work we presented a low cost, high performance, and straightforward mechanical ventilator that can be deployed in public health emergencies rapidly. We successfully mimicked the functionality of the two primary modes (*i.e*., CM and AM) of operation for modern ventilators. In addition, we achieved the accuracy and consistency of a certified commercialized ventilator. To further improve on the device, more modes of operation can be implemented easily as the underlying design and architecture accommodates for these alternative modes. In addition, sensor for monitoring the blood oxygen saturation level, exhale CO_2_ level, temperature, EKG, and heartbeat can be easily integrated.

## Materials and Methods

### Hardware architecture

The artificial respirator device is constructed of eight major components. The specifications and rationale behind choosing these components are explained in the following:

- *Stepper motor:* The maximum safe pressure that can be exerted on the respiratory system is 40 cmH_2_O. To generate this pressure with the glass syringe (plunger diameter of 82 mm), we need a force of 20.7 N. To deliver 1000 mL air, we need an arm length of 95 mm, which gives a torque requirement of 1.97 N⋅m for the motor. Considering that the pressure transmission is not always 100%, we chose a Nema 23, bipolar, 3 N⋅m stepper motor. Stepper motors with higher torque ratings can be utilized for higher respiration rates and *I*:*E* ratios.
- *Motor driver:* we chose a digital stepper driver with a peak current rating of 4.2A to drive the stepper motor with our microcontroller. The inputs of the driver are optically isolated from the microcontroller to prevent backpropagation of back EMF noise or any EM noise from the stepper motor. We used 400 pulse/rev setting to achieve a balance between smooth motion and stressing the microcontroller.
- *Crank-Shaft Linkage:* We made the crank-shaft linkage from the components illustrated in figure 1. The crank-shaft linkage components include a clevis rod end, a piece of high-strength steel threaded rod, a partially threaded rod end bolt, a ball joint linkage, medium-strength steel hex nut, a high-strength steel threaded rod, a piece aluminum sheet, and a flange mount shaft collar to connect the crank to the stepper motor. To join the top end of the glass syringe to the rod end bolt, we used a fast curing urethane resin that has similar performance and mechanical characteristics to ABS plastic. Figure 8 illustrates the procedure.
- *Glass Syringe:* We used a 1000 mL glass syringe manufactured by BLG. The syringe has a 15 mm opening at the tip, which is large enough not to slow down the system for fast actuation rates.
- *Power supply:* A 48V switching power supply with a current rating of 10A is used to power the stepper motor.
- *Microcontroller:* We employed an 84 MHz, 32-bit ARM core microcontroller.
- *Sensors:* A differential pressure sensor (Honeywell HSCDRRN160MDSA3) is used to monitor the pressure via the SPI interface.
- *Interface:* A 2.4-inch TFT color display was used for this work with 12 pushbutton switches and 2 limit switches. The pushbutton switches are soldered on a PCB board with no debounce capacitors. The limit switches are employed to enable homing and recalibration of the stepper motor’s position in each breathing cycle. Shield wires are used to eliminating EM interference of the stepper motor with the limit switch signals. For the alarm, we used an internally driven Piezo with an operating frequency of 4.1kHz (C8 on Piano).

**Figure 8.**
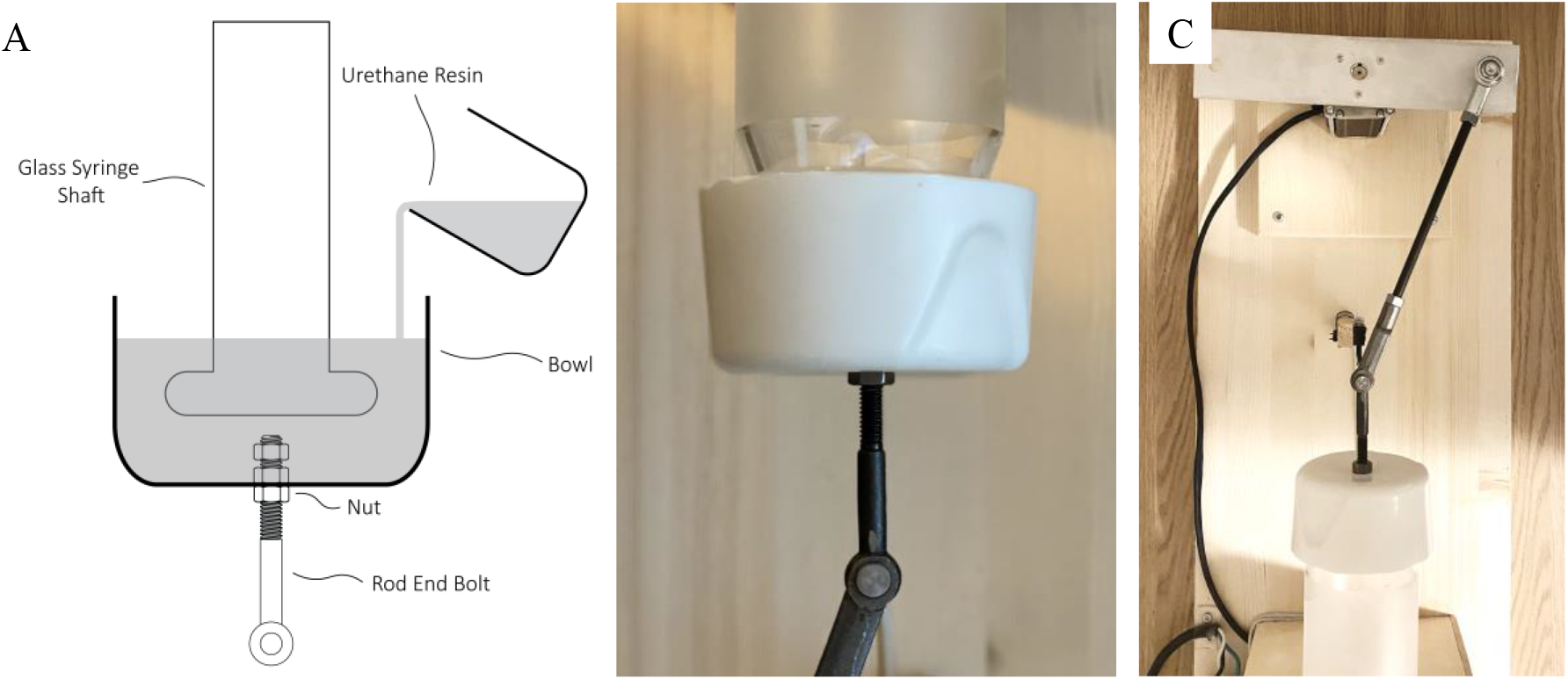
The procedure for attaching the rod end bolt to the top of the glass syringe plunger. (A) After the precursors of the urethane resin are mixed and degassed in a desiccator, it is poured to the bowl, which holds the rod end bolt. Two nuts are used inside the container to make it easy to unscrew the rod end bolt if needed. Items are not to scale on the drawing. (B) The result of the procedure in (A) showing the attachment. The diameter of the plunger is 82 mm. (C) Top view of the crank-shaft linkage connecting the stepper motor to the plunger of the glass syringe.

### Setup Design

We used pine lumber (3/4”, 12“×36”) for the chassis of the device and pine slabs and studs for mounting the glass syringe, limit switches and making the housing for the electronics. For the final commercialized device, these components will be hosted in an enclosure.

### Software architecture

We designed and implemented two similar state machines for the CM and AM. The following section describes the high-level logic of the state machines. Figure 9 illustrates the high-level logic of the CM and AM states. Both modes have the same architecture but with slightly different logic. Details of each state are described below:

**Figure 9.**
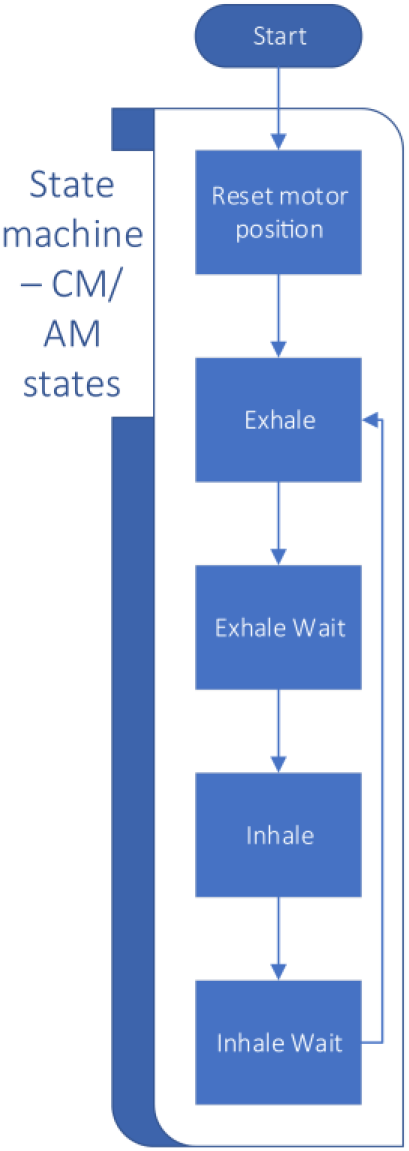
Software state diagram and flow.

- *Reset Motor State:* This state is only executed once after the transition from the ‘Idle’ state to the ‘CM/AM’ states. The goal of this state is to calculate all cycle parameters and move the motor clockwise (CW) to a known 180° position. This position is the same position in which the motor will be in at the end of the inhale cycle. The cycle parameters are calculated in this state. This includes the motor’s speed, final rotation steps, and breathing cycle wait times.
- *Exhale State:* This state starts the exhale cycle and moves the motor counterclockwise (CCW). This state ends once the stepper motor reaches the final rotation steps.
- *Exhale Wait State:* This state holds the motor in position for a set period. This period is determined at the start of the cycle from the *I*:*E* ratio, respiration rate, and the tidal volume parameters. In the AM, the exhale wait state has additional logic to monitor pressure to detect the patient’s attempt to inhale.
- *Inhale State:* Similar to the reset state, the motor is moved CW back to the 180° position in this state. The start of the inhale state is also considered the beginning of a new breathing cycle. Therefore, any setting parameters such as respiration rate, *I*:*E* ratio, and tidal volume will be updated.
- *Inhale Wait State:* Similar to the Exhale Wait state, the motor is held in position for a set period of time. This state also calculates and displays the maximum flow rate, peak pressure, PEEP, and plateau pressure.

The breathing cycle is then repeated until the device is stopped, rebooted, or powered off.

### Cost analysis

The list of the materials and the cost to build a prototype of the device is tabulated in table 2. The cost amount per component displayed in this table is cost for 1 unit of each component. The total bill of material (BOM) can be significantly reduced if purchased and assembled in larger quantities. Based on estimated quotes we received from the suppliers, if the components are purchased in units of 1000, the total BOM per device drops to approximately $250/device.

**Table 2.** Bill of Material for the ventilator.

| Components | Numbers Used | Unit Price (USD) |
| --- | --- | --- |
| Power supply | 1 | \$ 14.27 |
| Stepper Motor | 1 | \$ 33.75 |
| Motor Driver | 1 | \$ 23.64 |
| 1000 mL Glass Syringe | 1 | \$ 200.50 |
| Microcontroller | 1 | \$ 15.99 |
| LCD | 1 | \$ 2.50 |
| Pushbuttons | 12 | \$ 1.79 |
| Limit Switches | 2 | \$ 4.97 |
| Buzzer | 1 | \$ 6.09 |
| Rod end bolt | 1 | \$ 6.32 |
| Clevis Rod Ends | 1 | \$ 5.15 |
| Ball Joint Linkage | 1 | \$ 5.21 |
| 1ft High-Strength Steel Threaded Rod | 0.5 | \$ 9.33 |
| Pressure Sensor | 1 | \$ 35.70 |
| Shaft Collar | 1 | \$ 2.00 |
| Miscellaneous (wood, bolts, screws, etc.) | | \$ 20.00 |
| **Total** | | **\$ 387.21** |

## Acknowledgment

The author benefited substantially from Ms. Mahdieh Chavoshi (Nurse Anesthetist) and Ms. Farnoush Mirvakili, MD on the clinical aspect of ventilators and our design described in this work.

## Author Contributions

S.M.M. conceived the design, built the hardware, and performed all the experiments. D.S. designed and implemented the software. S.M.M. and D.S. performed the data analysis and wrote the manuscript. R.L. provided supervision and guidance. All authors read and commented on the manuscript.

## Data Summary for Test Cases

We performed 13 experiments with the test lung to evaluate the performance of the APA ventilator for various scenarios. The results are plotted in Figure 4. The values for the set parameter and measured data for each cycle are tabulated in Table S1.

**Table S1.** Result summary for different setting configurations. Each experiment is performed for 10 consecutive cycles.

| Test No. | Set Parameters |  |  |  |  | Measured Values |  |  |
| --- | --- | --- | --- | --- | --- | --- | --- | --- |
| | $VT$ (mL) | $RR$ (b/min) | $I:E$ ratio | $End$-inspiratory hold (%) | $PEEP$ (cmH<sub>2</sub>O) | $P_{peak}$ (cmH<sub>2</sub>O) | $P_{plateau}$ (cmH<sub>2</sub>O) | $F_{max}$ (L/min) |
| 1 | 750 | 10 | 1:2 | 50 | 5.22± 0.23 | 41.53±0.44 | 37.69± 0.64 | 58.61± 0.02 |
| 2 | 550 | 18 | 1:2 | 50 | 5.64±0.13 | 33.24± 0.66 | 22.98±0.54 | 82.00±0.08 |
| 3 | 400 | 24 | 1:2 | 50 | 5.92±0.07 | 28.00±0.41 | 21.11±0.06 | 85.46 |
| 4 | 300 | 30 | 1:2 | 50 | 6.04±0.04 | 25.16±1.61 | 19.01±0.25 | 81.92±0.14 |
| 5 | 550 | 10 | 1:2 | 50 | 5.45±0.15 | 24.78±0.23 | 22.01±0.21 | 45.59±0.01 |
| 6 | 550 | 12 | 1:2 | 50 | 5.60±0.13 | 26.31±0.22 | 22.5±0.23 | 54.70±0.02 |
| 7 | 550 | 14 | 1:2 | 50 | 5.63±0.12 | 28.14±0.42 | 22.92±0.36 | 63.82±0.02 |
| 8 | 550 | 16 | 1:2 | 50 | 5.69±0.11 | 30.64±0.61 | 23.18±0.50 | 72.92±0.04 |
| 9 | 500 | 13 | 1:1 | 50 | 5.75±0.08 | 23.11±0.27 | 20.70±0.03 | 36.58±0.01 |
| 10 | 500 | 13 | 1:2 | 50 | 5.61±0.08 | 25.20±0.34 | 20.95±0.07 | 54.83±0.01 |
| 11 | 500 | 13 | 1:3 | 50 | 5.54±0.16 | 28.59±0.40 | 21.10±0.11 | 73.06±0.05 |
| 12 | 500 | 13 | 1:4 | 50 | 5.30±0.13 | 33.87±0.27 | 21.10±0.52 | 91.37±0.05 |
| 13 | 500 | 13 | 1:5 | 50 | 5.20±0.13 | 39.52±1.11 | 20.95±0.91 | 109.45±0.10 |

For experiments 1 to 4, we kept the *I*:*E* ratio constant at 1:2 but increased the *RR* from 12 b/min to 30 b/min while decreasing the *VT* from 750 mL to 300 mL. As we decreased the tidal volume, both the peak pressure and plateau pressure decreased as well. This correlation can be explained by the fact that less air is moved to the test lung; therefore, a smaller pressure is developed inside it. For experiments 5 to 8, we kept the tidal volume and *I*:*E* ratio constant and increased the respiration rate. As expected, the flow rate increases as well as the difference between the peak pressure and plateau pressure increases. To show this effect in more depth, for experiments 9 to 13, we kept the *VT* at 500 mL and *RR* at 13 b/min but increased the *I*:*E* ratio from 1:1 to 1:5. Similar to the previous case, since the volume is kept constant, the plateau pressure is also constant; however, the peak pressure increases, which can be explained by the fact the resistive pressure is increasing due to the increase in flow rate. More details are provided in the next section. This behavior of the test lung is similar to that of the human lungs.

## Dynamics of the Respiratory System

The lung is a porous structure made of soft tissues and a complex network of airway branches. Due to the inhomogeneity of its structure, the pressure developed in the lungs has three components:

- *Resistive Pressure (P_resistive_)* which is the contribution from the resistive properties of the respiratory system. Viscous and turbulent losses related to flow of gas through the airway tree and the deformation of parenchymal and chest wall tissues are probably the main contributors to the resistive pressure (*1*). At small flow rates, the resistive pressure is a linear function of the flow rate 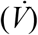. However, at higher flow rates (*e.g.*, low *I*:*E* ratios, exercising), the resistive pressure scales nonlinearly, often in the form of 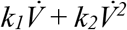, where *k_1_* and *k_2_* are determined empirically.
- *Elastic Pressure (P_elastic_)* which is the contribution from the recoil of the lungs and chest wall to their relaxed states when inflated with air either by contraction of the diaphragm and intercostal muscles or by mechanical insufflation (*e.g.*, a ventilator). Therefore, the elastance of the respiratory system (*E_rs_*) can be seen as a linear summation of the elastance from the chest walls (*E_cw_*) and the elastance from the lungs (*E_l_*). The reciprocal of the respiratory system’s elastance gives its compliance (*C_rs_*). The elastic pressure is a function of the tidal volume.
- *Inertial Pressure (P_inertial_)* which is the contribution from the inertial forces from the chest wall, airway tree, and parenchymal tissues. The inertial pressure is a function of volume’s acceleration 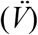.

Therefore, the total pressure is (*2*, *3*):

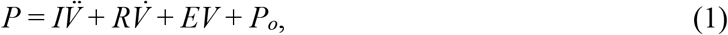

where *I*, *R*, and *E* are the internal, elastic, and resistive properties of the respiratory system. The *P_o_* is the distending pressure at the end of expiration.

The resistance (*R*) can be estimated from the difference between the peak pressure and plateau pressure divided by the flow rate. Any alterations in *P_resistive_* (for a specified flow rate) can reflect changes in the airway caliber (*4*).

At zero flow rate, the elastance of the respiratory system can be determined from the difference between the plateau pressure and the *PEEP* divided by the tidal volume. It is important to note that the dynamic elastance is higher than static elastance in general, which is due to the viscoelasticity and gas redistribution. Figure S1 illustrates the contribution of each component of the pressure in the respiratory system.

**Figure S1.**
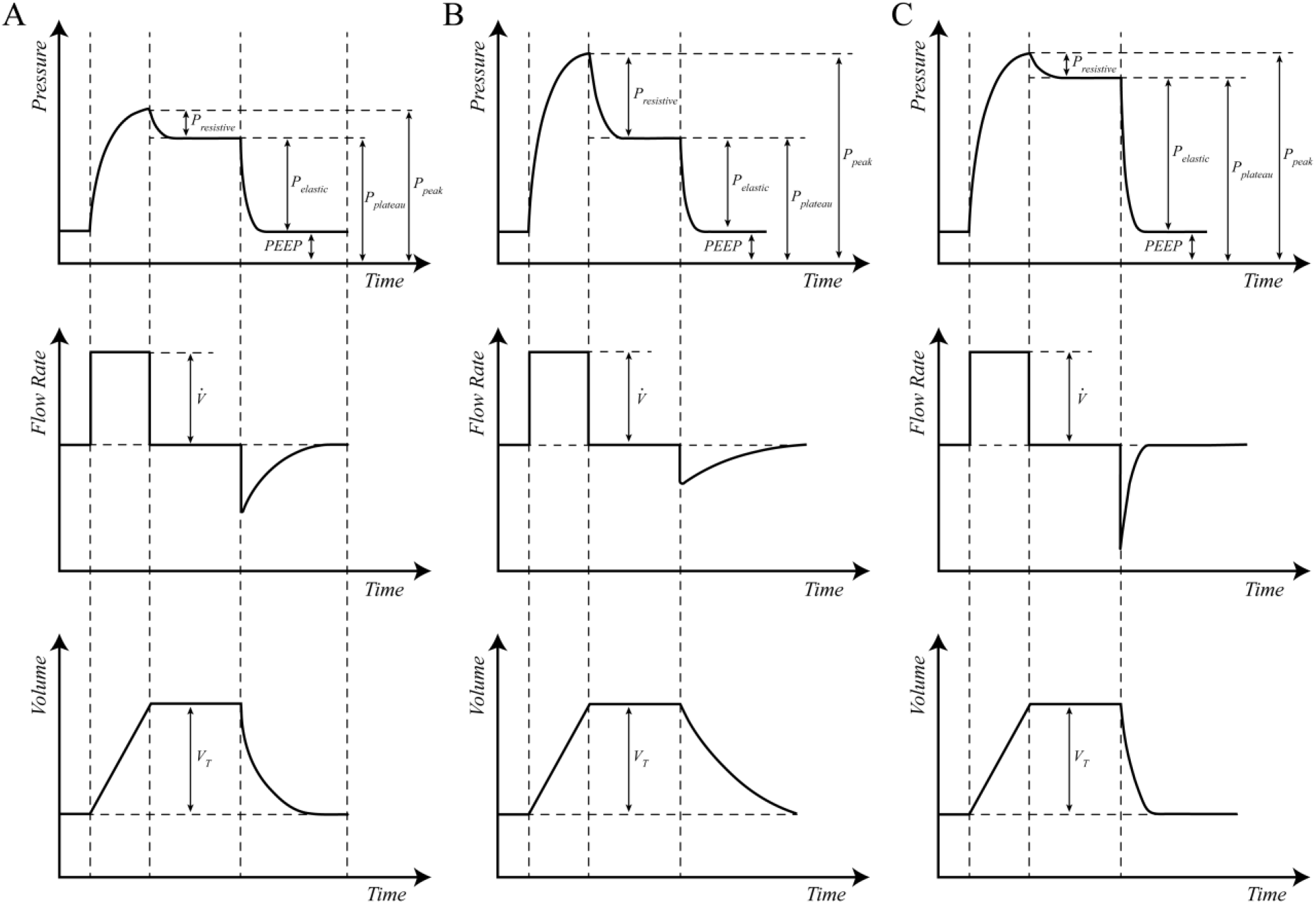
Pressure, flow rate, and volume profiles in volume-controlled ventilation (at constant flow rate). (A) Pressure, flow rate, and volume profiles for a healthy patient. In this the case the resistive pressure has a small contribution to the peak pressure. While the elastic pressure has the largest contribution. (B) The peak pressure is increased while the plateau remains similar to that of in (A). This indicates an increase in the resistive pressure. This condition occurs in cases such as bronchospasm, mucous plug, retained secretions, and ETT tip occlusion. (C) The elastic pressure is increased in this case, while the resistive pressure contribution is similar to that of in part (A). Conditions such as ARDS, pneumonia, pneumothorax, and pulmonary oedema lead to an increase in the elastic pressure.

## Actuator Kinematics

The position of the plunger is almost proportional to the cosine of the angle between the crank axis and axis of the plunger (Figure S2). From the cosine law, we can find the position of the plunger (*s*) as a function of the crank radius (*r*) – which is half of the linear stroke, and length of the connecting rod (*l*) as the following equation suggests:

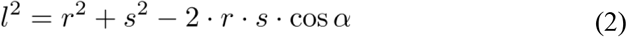

**Figure S2.**
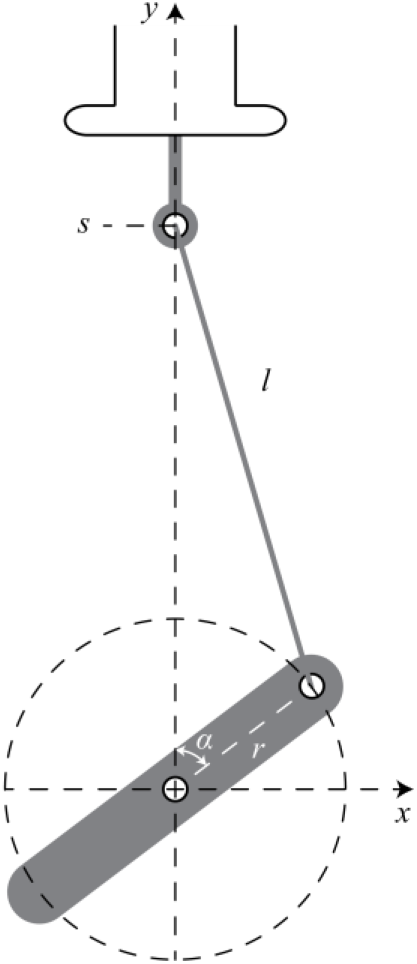
Diagram of the geometry of the crank-shaft linkage.

The position of the plunger corresponds to the volume that the syringe can deliver. By rearranging the terms in equation 2, we find the *s* to be:

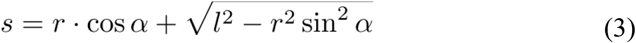

Therefore, the tidal volume (*VT*) as a function of rotation stroke (*α_i_* - *α_f_*) can be found to be:

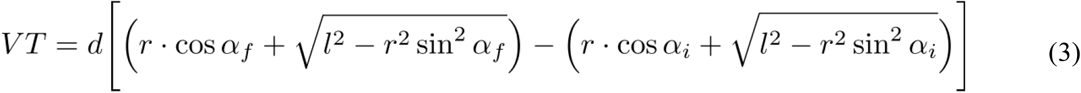

where *d* is the volume capacity of the glass syringe (*i.e.*, 1000 mL) over the length the plunger should travel inside the barrel to achieve that volume capacity (*e.g.*, the amplitude of the maximum linear stroke). For extreme cases of *α* = 0° and *α* = 180° we obtain *VT* = 2*rd*. Since *r* is half of the maximum linear stroke, we obtain *VT* = 1000 mL as expected.

